# Tradict enables accurate prediction of eukaryotic transcriptional states from 100 marker genes

**DOI:** 10.1101/060111

**Authors:** Surojit Biswas, Konstantin Kerner, Paulo José Pereira Lima Teixeira, Jeffery L. Dangl, Vladimir Jojic, Philip A. Wigge

**Author notes:** Correspondence, [† Contributed equally].

## Abstract

Transcript levels are a critical determinant of the proteome and hence cellular function. Because the transcriptome is an outcome of the interactions between genes and their products, it may be accurately represented by a subset of transcript abundances. We developed a method, Tradict (<u>tra</u>nscriptome pre<u>dict</u>), capable of learning and using the expression measurements of a small subset of 100 marker genes to predict transcriptome-wide gene abundances and the expression of a comprehensive, but interpretable list of transcriptional programs that represent the major biological processes and pathways of the cell. By analyzing over 23,000 publicly available RNA-Seq datasets, we show that Tradict is robust to noise and accurate. Coupled with targeted RNA sequencing, Tradict may therefore enable simultaneous transcriptome-wide screening and mechanistic investigation at large scales.

## Introduction

As the critical determinant of the proteome and therefore cellular status, the transcriptome represents a key node of regulation for all life^1^. Transcriptional control is managed by a finely tuned network of transcription factors that integrate environmental and developmental cues in order to actuate the appropriate responses in gene expression^2–4^. Importantly, the transcriptomic state space is constrained. Pareto efficiency constraints suggest that no gene expression profile or phenotype can be optimal for all tasks, and consequently, that some expression profiles or phenotypes must come at the expense of others^5,6^. Furthermore, across all major studied kingdoms of life, cellular networks demonstrate remarkably conserved scale-free properties that are topologically characterized by a small minority of highly connected regulatory nodes that link the remaining majority of sparsely connected nodes to the network^7–9^. These theories suggest that the effective dimension of the transcriptome should be far less than the total number of genes it contains. If true to a large enough extent, it may be possible to faithfully compress and prospectively summarize entire transcriptomes by measuring only a small, carefully chosen subset of it.

Indeed, previous studies have exploited this reduced dimensionality to perform gene expression imputation for microarray data for missing or corrupted values^10–12^. Others have extended these intuitions to predict expression from probe sets containing a few hundred genes^13,14^. However, prediction accuracies have been modest and usually limited to 4,000 target probes/genes. Recently, several studies examined the transcriptomic information recoverable by shallow sequencing especially as it applies to single-cell experiments^15–18^. Jaitin *et al.* (2014) and Pollen *et al.* (2014) demonstrated that only tens of thousands of reads are required per cell to learn and classify cell types *ab initio*^16,18^. Heimberg *et al.* (2016) extended these findings and demonstrated that the major principal components of a typically sequenced mouse bulk or single-cell expression dataset may be estimated with little error at even 1% of the depth^15^. Though these approaches advance the notion of strategic transcriptome undersampling, they only recover broad transcriptional states and are restricted to measuring only the most abundant genes. During sample preparation -- arguably the most expensive cost of a multiplexed sequencing experiment -- shallow sequencing-based approaches still utilize protocols meant for sampling the entire transcriptome and therefore consume more resources than necessary. Furthermore, given that the expression of even the most abundant genes is highly skewed, sequencing effort is wastefully distributed compared to an approach that chooses which genes to measure more wisely. Finally, it is still not clear from sample sizes and biological contexts previously studied whether the low dimensionality of the transcriptome may be leveraged unconditionally (or nearly so) across organism and application.

In this work, we introduce Tradict (<u>tra</u>nscriptome pre<u>dict</u>), a robust-to-noise and probabilistically sound algorithm, for inferring gene abundances transcriptome-wide, and predicting the expression of a transcriptomically comprehensive, but interpretable list of *transcriptional programs* that represent the major biological processes and pathways of the cell. Tradict makes its predictions using only the expression measurement of a single, context-independent, machine-learned subset of 100 marker genes. Importantly, Tradict’s predictions are formulated as posterior distributions over unmeasured genes and programs, and therefore simultaneously provide point and credible interval estimates over predicted expression. Using a representative sampling of over 23,000 publicly available, transcriptome-wide RNA-Seq datasets for *Arabidopsis thaliana* and *Mus musculus,* we show Tradict prospectively models program expression with striking accuracy. Our work demonstrates the development and large-scale application of a probabilistically reasonable multivariate count/non-negative data model, and highlights the power of directly modeling the expression of a comprehensive list of transcriptional programs in a supervised manner. Consequently, we believe that Tradict, coupled with targeted RNA sequencing^19–24^, can rapidly illuminate biological mechanism and improve the time and cost of performing large forward genetic, breeding, or chemogenomic screens.

## Results

### Assembly of a comprehensive training collection of transcriptomes

We downloaded all available Illumina sequenced publicly deposited RNA-Seq samples (transcriptomes) on NCBI’s Sequence Read Archive (SRA). Among samples with at least 4 million reads, we successfully downloaded and quantified the raw sequence data of 3,621 and 27,450 transcriptomes for *A. thaliana* and *M. musculus,* respectively. After stringent quality filtering, we retained 2,597 (71.7%) and 20,847 (76.0%) transcriptomes comprising 225 and 732 unique SRA submissions for *A. thaliana* and *M. musculus,* respectively. An SRA ‘submission’ consists of multiple, experimentally linked samples submitted concurrently by an individual or lab. We defined 21,277 (*A. thaliana*) and 21,176 (*M. musculus*) measurable genes with reproducibly detectable expression given our tolerated minimum sequencing depth and mapping rates (see Supplemental Information “Materials and Methods” for further information regarding data acquisition, transcript quantification, quality filtering, and expression filtering). We hereafter refer to the collection of quality and expression filtered transcriptomes as our *training transcriptome collection*.

In order to assess the quality and comprehensiveness of our training collection, we performed a deep characterization of the expression spaced spanned by these transcriptomes. We found that the transcriptome of both organisms was highly compressible and that the primary drivers of variation were tissue and developmental stage (Figure 1a-b, Figure S1), with many biologically realistic trends within each cluster (Supplemental Analysis 1). We additionally examined the distribution of submissions across the expression space, compared inter-submission variability within and between tissues, inspected expression correlations among genes with well-established regulatory relationships, and assessed the evolution of the expression space across time. These investigations revealed our training collection is of high and reproducible technical quality, reflective of known biology, and stable (Supplemental Analysis 1, Figures S2-S4). Given additionally the diversity of tissues, genetic perturbations, and environmental stimuli represented in the SRA, these results, taken together, suggest that our training collection is an accurate and representative sampling of the transcriptomic state space that is of experimental interest for both organisms.

**Figure 1.**
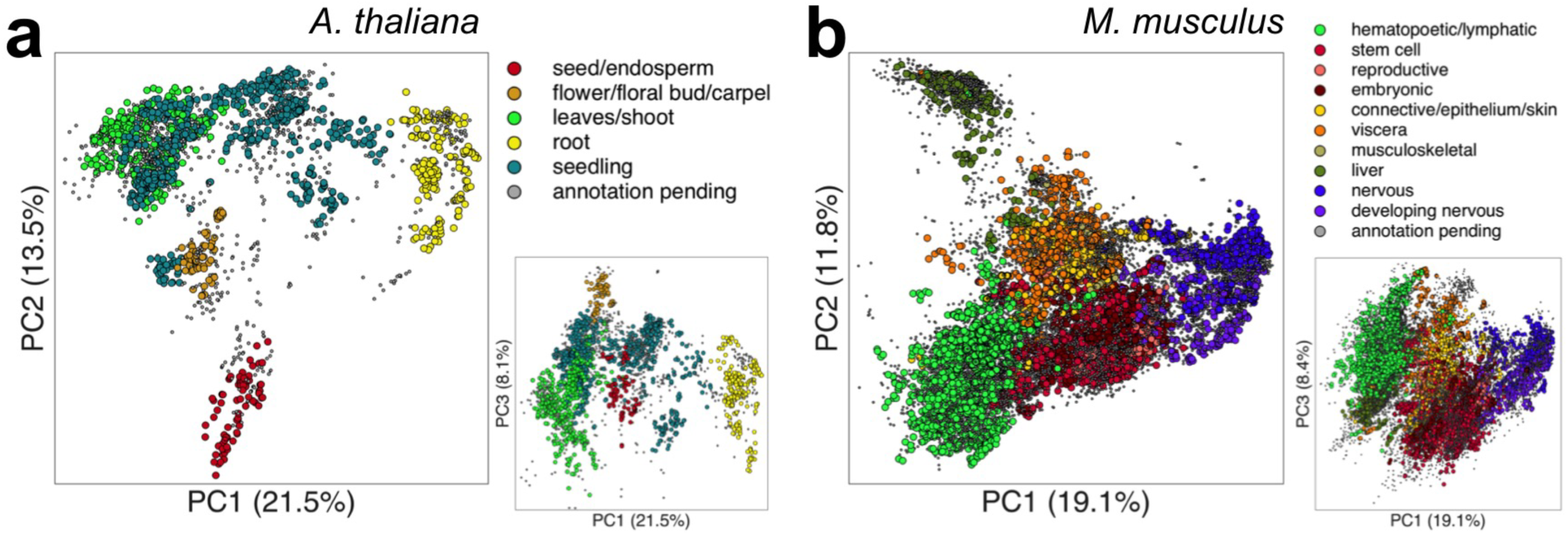
The primary drivers of variation in our training transcriptome collection are developmental stage and tissue. a)*A. thaliana,* b) *M. musculus.* Also shown are plots of PC3 vs. PC1 to provide another perspective.

### Tradict - algorithm overview

Given a training transcriptome collection, Tradict encodes the transcriptome into a single subset of globally representative *marker* genes and learns their predictive relationship to the expression of a comprehensive collection of transcriptional programs (e.g. pathways, biological processes) and to the rest of the genes in the transcriptome. Tradict’s key innovation lies in using a Multivariate Normal Continuous-Poisson (MVN-CP) hierarchical model to model marker latent abundances -- rather than their measured, noisy abundances -- jointly with the expression of transcriptional programs and the abundances of the remaining non-marker genes in the transcriptome. In so doing, Tradict is able to 1) efficiently capture covariance structure within the non-negative, right-skewed output typical of sequencing experiments, and 2) perform robust inference of gene set and non-marker expression even in the presence of significant noise.

Figure 2 illustrates Tradict’s general workflow. Estimates of expression are noisy, especially for low to moderately expressed genes. Given samples are often explored unevenly and that the *a priori* abundance of each gene differs, the level of noise in a gene’s measured expression for a given sample varies, but it can be estimated. Therefore, during training, Tradict first learns the log-latent, denoised abundances for each gene in every sample in the training collection using the lag transformation^25^. It then collapses this latent transcriptome into a collection of predefined, comprehensive collection of *transcriptional programs* that represent the major processes and pathways of the cell related growth, development, and response to the environment (Supplemental Tables 3-4). In this work, we focus on creating a Gene Ontology derived panel of transcriptional programs, in which the first principal component of all genes contained within an appropriately sized and representative GO term is used to define an accordingly named transcriptional program^26,27^. The expression values of these programs are then encoded using an adapted version of the Simultaneous Orthogonal Matching Pursuit algorithm into a small subset of marker genes selected from the transcriptome^28,29^. Tradict finally stores the mean and covariance relationships between the log-latent expression of the selected markers, the transcriptional programs, and the log-latent expression of the remaining non-marker genes at the Multivariate Normal layer of the MVN-CP hierarchical model (Figure 2a).

**Figure 2.**
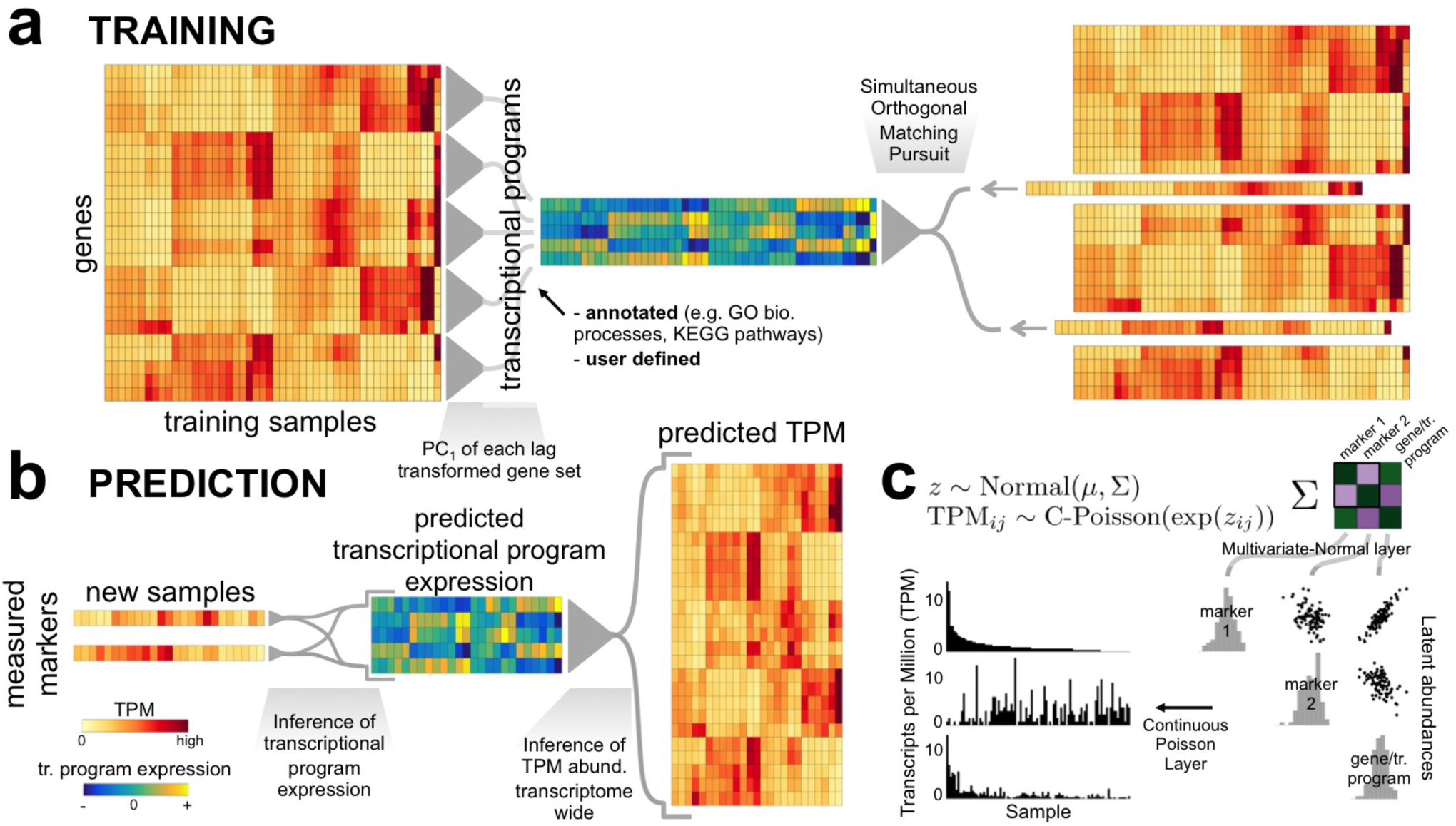
Tradict’s algorithmic workflow. a) During training, the transcriptome is first quantitatively summarized in terms of a collection of a few hundred, biologically comprehensive transcriptional programs. These are then decomposed into a subset of marker genes using an adaptation of the Simultaneous Orthogonal Matching Pursuit algorithm. A Multivariate Normal Continuous-Poisson hierarchical model is used as a predictive model to capture covariance relationships between markers, transcriptional programs, and all genes. b) During prediction, Tradict predicts the expression of transcriptional programs and all genes in the transcriptome using the expression measurements of the marker genes. c) The Multivariate Normal Continuous-Poisson hierarchy enables Tradict to efficiently model statistical coupling between the non-negative expression measurements typical of sequencing experiments. This is done by assuming that associated with each observed, noisy TPM measurement, there is an unmeasured (denoised), latent abundance the logarithm of which comes from a Multivariate Normal distribution over all genes and transcriptional programs.

Prospectively, only the expression of these marker genes needs to be measured and the expression of genes and/or transcriptional programs can be inferred as needed. During prediction, Tradict uses the observed marker measurements as well as their log-latent mean and covariance learned during training, to estimate -- via MCMC sampling -- the posterior distribution over the log-latent abundances of the markers^30^. Though a simply a consequence of proper inference of our model, this denoising step adds considerable robustness to Tradict’s predictions. From this estimate, Tradict uses covariance relationships learned during training to estimate the conditional posterior distributions over the remaining non-marker genes and transcriptional programs (Figure 2b). From these distributions, the user can derive point estimates (e.g. posterior mean or mode), as well as measures of confidence (e.g. credible intervals). A complete description of the entire algorithm can be found in the “Tradict algorithm” section of the Materials and Methods in the Supplemental Information.

### Tradict prospectively predicts gene and transcriptional program expression with superior and robust accuracy

To understand Tradict’s prospective predictive performance, we performed 20-fold cross validation on the training transcriptome collections for both *A. thaliana* and *M. musculus* and evaluated Pearson correlation coefficients (PCC) between predicted and actual expression for each fold when the remaining 95% of folds were used for training. To make this experiment as reflective of reality as possible, folds were divided by submission so that samples from the same set of experiments would not appear both in training and test sets. Because submissions to the SRA span a broad array of biological contexts, the total biological signal contained in any given test set likely exceeds that of what would be expected for typical application, which in turn would lead to overly optimistic estimates of prediction accuracy. We therefore evaluated *intra-submission accuracy*, in which PCC calculations are performed on ‘submission-adjusted’ expression values. To do this, for each gene and program, each test-set submission’s mean expression was subtracted from the expression values for all samples associated with that submission. In effect, this regresses out between-submission effects, and allows us to assess Tradict’s predictive performance one experiment at a time, as one would do in practice.

Figures 3a and 3c illustrate that the reconstruction performance for transcriptional programs in both organisms is strikingly accurate across all collected submissions. Quantitatively speaking, the average intra-submission PCCs for transcriptional programs are 0.94 and 0.93 for *A. thaliana* and *M. musculus,* respectively. This is despite lower prediction performance on gene expression (Figures 3b and 3d). Intuitively, this is because transcriptional programs are measured as linear combinations of the log-latent TPMs of the genes that comprise them, effectively smoothing over the orthogonal noise present in each gene’s expression prediction.

**Figure 3.**
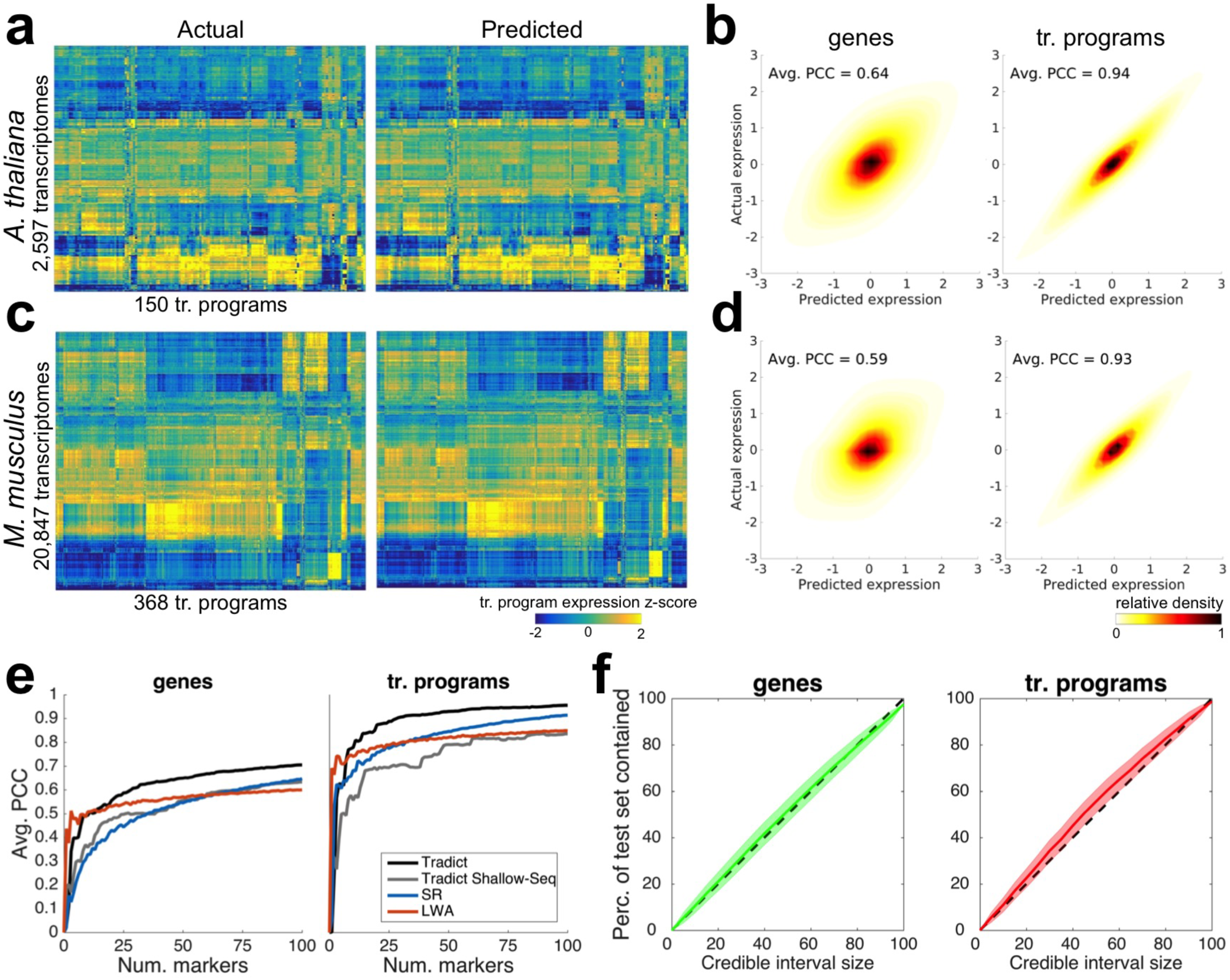
Tradict prospectively predicts gene and transcriptional program expression with superior and robust accuracy. Tradict’s prospective prediction accuracy during 20-fold cross validation of the entire training collection for both organisms. a) Heatmaps illustrating test-set reconstruction performance of all transcriptional programs for *A. thaliana.* Shown is the reconstruction performance for all samples in our transcriptome collection when they were in the test-set. b) Density plots of predicted vs. actual test-set expression for all genes (left) and transcriptional programs (right) for *A. thaliana,* after controlling for inter-sumbission biological signal. The intra-submission expression of each gene and transcriptional program was z-score transformed to make their expression comparable. c & d) Same as a & b, but for *M. musculus.* e) Comparison of Tradict’s performance vs. several baselines: SR (structured regression), LWA (locally weighted averaging), and Tradict Shallow-Seq. f) A posterior predictive check illustrating the concordance between Tradict’s posterior predictive distribution and the distribution of test-set expression values for genes (left) and transcriptional programs (right). Plotted is the percent of test-set observations contained within a credible interval vs. the size of the credible interval. A unique credible interval is derived for each gene/program. The “x=y” line is illustrated as a dotted black line. Shaded error bands depict the sampling distribution of this analysis across test-sets from a 20-fold cross validation on the *A. thaliana* dataset.

We also found Tradict’s performance to be superior to several baselines. These include two successful approaches developed in Donner *et al.* (2012) for microarray^14^, and a version of Tradict that uses the 100 most abundant genes as its selected markers (Figure 3e, Figure S5, Supplemental Analysis 2). The former baselines rely on structured regression (SR) and locally weighted averaging (LWA), linear/parametric and non-parametric methods, respectively. The latter baseline examines the utility of simple shallow sequencing by using the most abundant genes as markers for Tradict (see Supplemental Analysis 2 for a more detailed description). Figure 3e illustrates test-set intra-submission performance of each method as a function of the number of markers entered into the model. LWA demonstrates the quickest performance gain, but then saturates after 10 markers. This is likely because a non-linear kernel based approach makes the most efficient use of a few markers, but is adversely impacted by the curse of dimensionality as more markers are added. The parametric methods (Tradict, SR) navigate this dimensionality increase more efficiently and ultimately realize better performance for a still reasonable number of markers. Tradict outperforms SR and Tradict Shallow-Seq, ultimately obtaining a PCC between predicted and actual expression of 0.71 for genes and 0.96 for transcriptional programs. This suggests Tradict’s probabilistic framework is more reasonable than SR’s and that Tradict’s marker selection is more optimal than picking the most abundant genes. We additionally found Tradict’s predictions were robust to noise in the form of low sequencing depth and/or corrupt marker measurements (Figure S5 and S10, Supplemental Analysis 2 and 5), which we attribute to its probabilistic framework, in which training and prediction are performed in the space of denoised latent abundances.

To further assess the validity of Tradict’s modeling assumptions, we examined how Tradict’s posterior predictive distribution matched the distribution of test-set gene and program expression values. Specifically, we performed a posterior predictive check in which we asked what percent of test-set gene or program expression values fall within an X% credible interval, where a unique interval is defined for each gene/program^30^. If Tradict’s posterior predictive distribution is reasonable then X% of the true expression values should fall within this interval for any X. Figure 3f illustrates the results of this analysis as performed on disjoint test-sets from a 20-fold cross validation on the *A. thaliana* dataset. On average, the X% credible interval captures X% of test-set observations for any choice of X. The posterior predictive distribution for transcriptional programs may be slightly too wide at moderate interval sizes (30-70%), which would make Tradict more conservative (higher type II error rate) than it should be. However, in practice it is accurate (p-value = 0.24, *t*-test) for larger, more standard interval sizes (e.g. 95%). We conclude that Tradict’s probabilistic modeling assumptions capture unseen data well.

We next characterized Tradict’s limitations through error, power, program annotation robustness analyses, and a timing and memory analysis (Supplemental Analysis 3-4). These analyses revealed that training-set expression variance and mean abundance correlated positively with both program and gene expression prediction performance (Supplemental Analysis 3.I, Supplemental Tables 3-4). Combined with program size as another predictor, these variables could account for most of the error (60% of total variance) in program expression prediction. A power analysis revealed that for both *A. thaliana* and *M. musculus,* 1,000 samples -- comprising approximately 100 submissions -- is sufficient for optimum performance (Supplemental Analysis 3.II). An examination of how gene-to-program mis-annotation rates influence predictive performance revealed that program expression prediction perfromance was robust up to a 20% mis-annotation rate and that gene expression prediction performance was completely robust to any level of mis-annotation. The latter result is a consequence of a statistical decoupling between gene and program expression prediction (Supplemental Analysis 3.III). Finally, Tradict’s training time and peak memory requirements scaled linearly with training set size, and increased 0.26 seconds and 1.1 Mb per sample. Prediction time was limited by MCMC sampling from the conditional posterior distributions of gene and program expression, and required 3.1 seconds per sample on average (Supplemental Analysis 4). Taken together, we conclude that the causes of Tradict’s errors are well understood and intuitive, and that Tradict’s sample requirements are reasonable, especially for major model organisms (Supplemental Table 5, Supplemental Analysis 3.II). Furthermore, Tradict is robust to noisily defined transcriptional programs, and its computational requirements scale well to large datasets.

### Case studies reveal the power of predicting and studying transcriptional program expression

To demonstrate how Tradict may be applied in practice, we focused on two case studies related to innate immune signaling -- one performed using bulk *A. thaliana* seedlings (detailed below), and the other using primary immune *M. musculus* cell lines (detailed in Supplemental Analysis 5; Figure S10). We trained Tradict on our full collection of training transcriptomes for each organism to produce two organism-specific Tradict models. Each was based on the selection of 100 markers learned from the full training transcriptome collection (Supplemental Tables 7-8) that we assert are globally representative, and context-independent. The case study samples do not, of course, appear in the collection of training transcriptomes.

### Tradict accurately predicts temporal expression patterns for a diverse panel of *A. thaliana* immune signaling mutants under different hormone perturbations

The hormones salicylic acid (SA) and jasmonic acid (JA) play a major, predominantly antagonistic regulatory role in the activation of plant defense responses to pathogens. Yang *et al.* (2016) investigated the effect of a transgenically expressed bacterial effector, HopBB1, on immune signaling in *A. thaliana*^31^. In their study, they performed a time course experiment, treating plants with MeJA (a JA response inducer), BTH (an SA mimic and SA response inducer), or mock buffer and monitored the transcriptome of bulk seedlings at 0 hr, 1 hr, 5 hr, and 8 hr post treatment. These experiments included several immune signaling mutants with differing degrees of response efficiency to MeJA and BTH treatment. Among other findings, they conclude that HopBB1 enhances the JA response, thereby repressing the SA response and facilitating biotrophic pathogen infection.

We asked to what extent strategic undersampling of the transcriptome and application of Tradict could quantitatively recapitulate the findings of Yang *et al.* (2016). Given Tradict’s near perfect accuracy on predicting the expression of transcriptional programs, we took a top down, but hypothesis driven approach to our analysis which first examined the expression of all transcriptional programs. Figure 4a illustrates the actual and predicted expression of all transcriptional programs in *A. thaliana* as a function of time and treatment. Here, Tradict reconstructs the expression of all transcriptional programs with an average PCC of 0.91.

**Figure 4.**
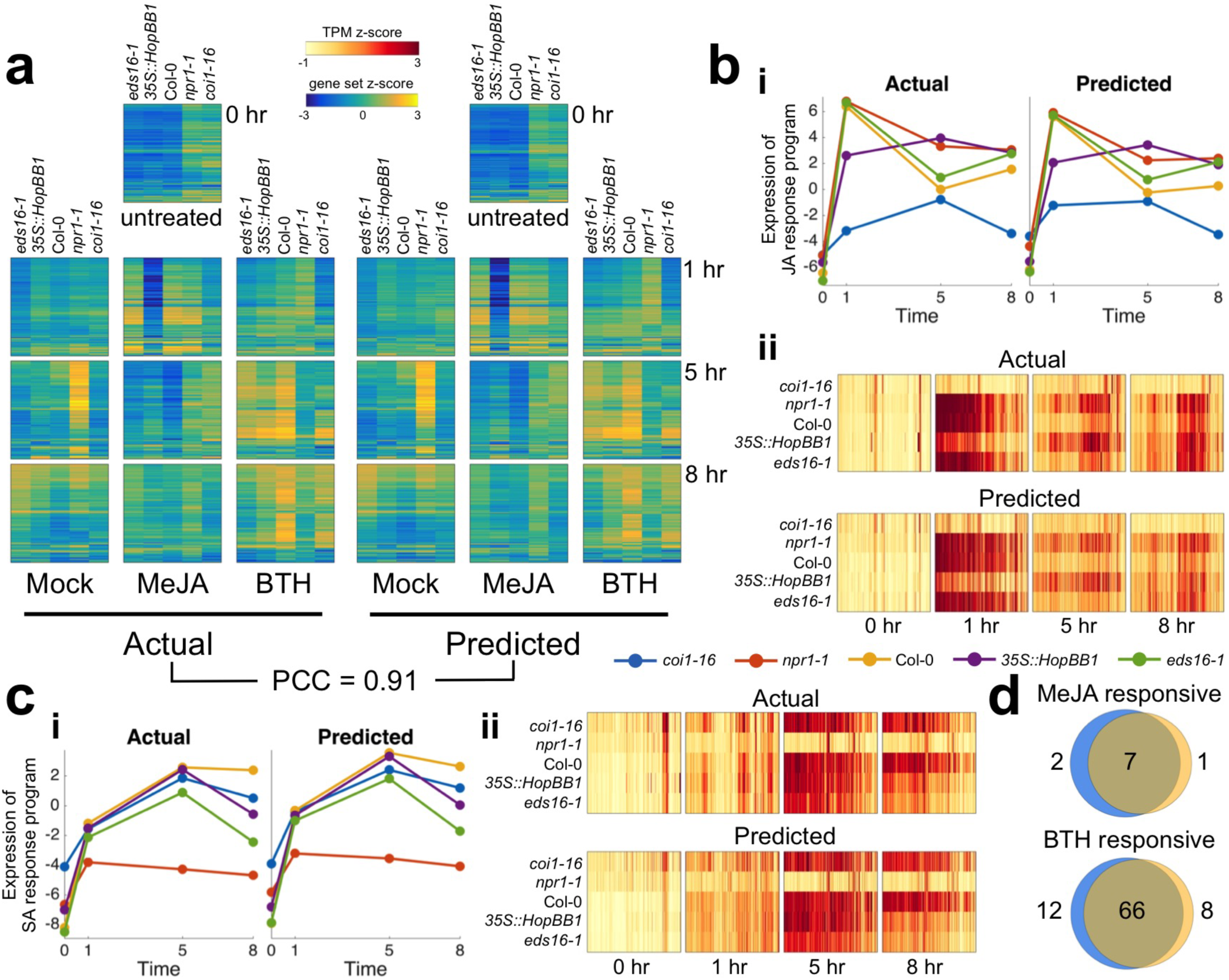
Tradict accurately predicts transcriptional responses across time in response to hormone perturbation in an *A. thaliana* innate immune signaling dataset. After being trained on the full *A. thaliana* training transcriptome collection, the selected set of 100 globally representative and context-independent markers were used to predict the expression of transcriptional programs and all genes for the transcriptomes presented in Yang *et al.* (2016). a) Actual vs. predicted heatmaps for the expression of all 150 transcriptional programs in *A. thaliana* across genotype, time, and hormone treatment. b) Predicted vs. actual expression of i) the JA response transcriptional program, and ii) the genes involved in the JA response program. c i-ii) Same as b, but for the SA response transcriptional program. d) Hypothesis free, differential transcriptional program expression analysis as performed on the actual expression of transcriptional programs vs those predicted by Tradict. Blue circles represent the actual and orange represent the predicted. All heatmaps are clustered in the same order across time, treatment, genotype, and between predicted and actual.

Recall that the genes participating in each of our transcriptional programs are pre-defined, in this work, by a carefully chosen, interpretable, but maximally representative set of GO biological processes. Therefore, given the goals of this study, we next examined the expression of the “response to jasmonic acid” and “response to salicylic acid” transcriptional programs. Figure 4b shows the expression behavior for the “response to jasmonic acid” transcriptional program across all the genotypes and time points upon MeJA treatment. More specifically, part (i) shows that the predicted expression and actual expression are qualitatively and quantitatively in agreement, both in magnitude and rank across the different genotypes. For example, as expected, *coi1*-16, which cannot sense JA, does not respond to the MeJA stimulus, while wildtype Col-0 does. However, even more subtle expression dynamics are captured by Tradict’s predictions. For example, *eds16-1* and *npr1-1* -- slightly and strongly impaired SA responders, respectively -- are slightly and strongly hyper-responsive to MeJA, respectively -- just as expected from the JA-SA antagonism. Furthermore, as demonstrated in Yang *et al.* (2016), the *35S::HopBB1* transgenic line exhibits a prolonged and sustained JA response for both the actual and predicted expression for this transcriptional program. Part (ii) of Figure 4b illustrates the expression of all the MeJA responsive genes in this transcriptional program. Again Tradict’s predictions are in good agreement with actuality, achieving a PCC of 0.72, and it’s visually clear that the expression magnitude of these genes positively correlates with the registered expression magnitude of the “response to jasmonic acid” transcriptional program. Figure 4c parts (i) and (ii) are presented in the same light as Figure 4b, but are instead illustrated for the SA response transcriptional program and constituent genes under BTH treatment. Again predictions match actuality, and the observed trends make biological sense^32^.

In order to illustrate Tradict’s use in hypothesis-free investigation, we performed a differential transcriptional program expression analysis for transcriptional programs affected by MeJA or BTH treatment (Figure 4d, see Methods). Differentially expressed transcriptional programs based on Tradict’s predictions versus actual measurements were highly concordant and biologically reasonable. Transcriptional programs differentially expressed with respect to MeJA treatment included “response to bacterium,” “defense response to fungus”, “response to wounding,” and “response to jasmonic acid” as expected. Transcriptional programs differentially expressed with respect to BTH treatment included various abiotic stress responses, “defense response to fungus”, “response to jasmonic acid” (via antagonism), and “response to salicylic acid,” again, as expected.

## Discussion

Tradict is an accurate, robust-to-noise method for predicting the expression of a comprehensive, but interpretable list of transcriptional programs that represent the major biological processes and pathways of the cell. Given the comprehensiveness, stability, and exponentially growing size of the training datasets we have assembled from publicly available sources, and as evidenced by our cross validation experiments, which were performed over thousands of samples deposited over the span of six years, the 100 markers Tradict learns are likely to be predictive independent of most contexts and applications. As illustrated through our case studies, examining the expression of these predicted transcriptional programs makes intuitive sense and provides a neat summary of underlying gene expression patterns.

Tradict additionally provides expression predictions for all genes in the transcriptome. Though its current gene expression prediction accuracy is less than ideal for more sensitive applications, Tradict’s performance is superior to previous efforts and is improving logarithmically in the number of samples. We attribute Tradict’s performance gains over previous methods first to improved measurement technology. Previous methods were developed for microarray, a substantially more noisy technology than RNA-Sequencing^10–14^. Consequently, training efficiency was lower as well as measurement accuracy of true expression, thus leading to modest prediction accuracy. By contrast, Tradict is meant to interface with sequencing based readouts of gene expression, a data type that is popular and proliferating exponentially as the time and price of sequencing continues to fall. Second, we believe Tradict’s probabilistic framework goes a step beyond previous efforts by modeling marker-gene and marker-program relationships not at the level of measured abundances, which are noisy, but at the level of latent abundances. Working in this denoised space naturally improves accuracy and affords robustness.

Taken together, we believe that Tradict coupled with target RNA sequencing can enable transcriptome-wide screening cheaply and at scale. Well-established commercial^19,20^ and non-commercial^21,33^ methods exist for targeted RNA sequencing in a multiplexed manner, and they are able to measure the expression of 10’s-100’s of genes with accuracy, making their use immediately compatible with Tradict. One method in particular, RASL-Seq^22–24^, does so cheaply with high precision and multiplexibility by directly probing total RNA and making efficient use of dual-indexing. We estimate that Tradict coupled with a time and resource efficient targeted RNA-sequencing protocol such as RASL-Seq could bring the cost of obtaining actionable transcriptome-wide information simultaneously for thousands to tens of thousands of samples to close to $1 per sample.

This scale could greatly benefit high-throughput breeding and screening applications. Forward genetic screens in most eukyarotic organisms require assaying 10^3^-10^4^ mutants. Small molecule, or more generally chemogenomic, drug screens often require screening thousands of molecules against multiple cell lines in multiple conditions. Agricultural screens -- whether for breeding or field phenotyping -- also require measuring thousands of individuals. Though in these cases a screen is made cheap and scalable by monitoring an easily selectable phenotype, new phenotyping architectures must be developed and optimized for each new screen (e.g. reporter lines, imaging hardware/software). Given the ubiquity of RNA, a transcriptome-wide screening approach would not suffer from such a drawback. Furthermore, and more importantly, though quickly interpretable, the phenotype being screened for is usually a uni-dimensional datum that offers little immediate insight into mechanism. In contrast, using Tradict to help perform transcriptome-wide screening could couple the process of hypothesis generation and mechanistic investigation. Here, we argue that the scalable monitoring of the expression of a comprehensive list of just a few hundred transcriptional programs affords an attractive balance of nuance and interpretability. Consequently, this efficient investigation, largely facilitated by Tradict, could greatly accelerate the pace of genetic dissection, breeding, and drug discovery.

## Acknowledgements

We thank Brian Cleary, Aviv Regev, Amaro Taylor-Weiner, Craig Bohrson, and Derek Lundberg for valuable discussions in preparing this manuscript. SB was supported by a Churchill Scholarship from the Winston Churchill Foundation of the United States and by a training grant from the NHGRI/NIH (T32 HG002295). PJPLT was supported by a fellowship from the Pew Latin American Fellows Program in the Biomedical Sciences. This work was additionally funded, in part, by a fellowship to PW from the Gatsby Foundation (GAT2373/GLB) and by grants to JLD from the National Institutes of Health (1RO1 GM107444), the Gordon and Betty Moore Foundation (GBMF3030), and the HHMI. JLD is an Investigator of the Howard Hughes Medical Institute.

## Code availability

A MATLAB implementation of Tradict is available at https://github.com/surgebiswas/tradict. All code to perform data downloads, analysis, and generate figures are available at https://github.com/surgebiswas/transcriptome_compression. Note that in order to make Tradict open-source and reach a wider audience we are developing a user-friendly, unit-tested Python version of Tradict that will be made available before publication.

